# OprF impacts *Pseudomonas aeruginosa* biofilm matrix eDNA levels in a nutrient-dependent manner

**DOI:** 10.1101/2023.03.01.530729

**Authors:** Erin K. Cassin, Sophia A. Araujo-Hernandez, Dena S. Baughn, Melissa C. Londono, Daniela Q. Rodriguez, Boo Shan Tseng

**Affiliations:** School of Life Sciences, University of Nevada Las Vegas, Las Vegas, NV, USA

**Keywords:** OprF, biofilm matrix proteins, eDNA, nutrient-dependent, biofilm maintenance

## Abstract

The biofilm matrix is composed of exopolysaccharides, eDNA, membrane vesicles, and proteins. While proteomic analyses have identified numerous matrix proteins, their functions in the biofilm remain understudied compared to the other biofilm components. In the *Pseudomonas aeruginosa* biofilm, several studies have identified OprF as an abundant matrix protein and, more specifically, as a component of biofilm membrane vesicles. OprF is a major outer membrane porin of *P. aeruginosa* cells. However, current data describing the effects of OprF in the *P. aeruginosa* biofilm is limited. Here we identify a nutrient-dependent effect of OprF in static biofilms, whereby Δ*oprF* cells form significantly less biofilm than wild type when grown in media containing glucose or low sodium chloride concentrations. Interestingly, this biofilm defect occurs during late static biofilm formation and is not dependent on the production of PQS, which is responsible for outer membrane vesicle production. Furthermore, while biofilms lacking OprF contain approximately 60% less total biomass than those of wild type, the number of cells in these two biofilms is equivalent. We demonstrate that *P. aeruginosa* Δ*oprF* biofilms with reduced biofilm biomass contain less eDNA than wild-type biofilms. These results suggest that the nutrient-dependent effect of OprF is involved in the maintenance of mature *P. aeruginosa* biofilms by retaining eDNA in the matrix.

**IMPORTANCE:** Many pathogens form biofilms, which are bacterial communities encased in an extracellular matrix that protects them against antibacterial treatments. The roles of several matrix components of the opportunistic pathogen *Pseudomonas aeruginosa* have been characterized. However, the effects of *P. aeruginosa* matrix proteins remain understudied and are untapped potential targets for antibiofilm treatments. Here we describe a conditional effect of the abundant matrix protein OprF on late-stage *P. aeruginosa* biofilms. A Δ*oprF* strain formed significantly less biofilm in low sodium chloride or with glucose. Interestingly, the defective Δ*oprF* biofilms did not exhibit fewer resident cells but contained significantly less extracellular DNA (eDNA) than wild type. These results suggest that OprF is involved in matrix eDNA retention in mature biofilms.

## INTRODUCTION

Biofilms are aggregates of bacterial cells encased in a self-produced extracellular matrix. The matrix protects resident cells from external assaults and is composed of exopolysaccharides, extracellular DNA (eDNA), membrane vesicles, and proteins (1). Many studies have reported the effects of exopolysaccharides, eDNA, and membrane vesicles on biofilm function. However, relatively few have investigated the roles of biofilm matrix proteins, even though matrix proteins have been suggested to play many vital functions in the biofilm (2–3). Since the late 2000s, researchers have used proteomic approaches to identify biofilm matrix proteins and gain insight into their roles, including several studies in the model biofilm organism *Pseudomonas aeruginosa*. Four different studies have identified OprF as an abundant matrix protein (4–7). Additionally, homologs of *P. aeruginosa* OprF have been identified in biofilm matrices of other organisms (8–10).

Within the *P. aeruginosa* biofilm, two populations of OprF protein exist: cell-associated and matrix-associated. In its more established cell-associated role, OprF is an OmpA family member and the major non-specific porin in *P. aeruginosa*, where it facilitates diffusion across the outer membrane (11). Multiple studies have examined biofilm formation after deletion of *oprF* or *ompA*, which eliminates both the cell- and matrix-associated protein pools (12–14). However, the impact of OprF and its OmpA homologs on biofilm formation is somewhat conflicting and may depend on conditions, such as oxygen or nutrient availability (15). One study shows that under aerobic conditions, a *P. aeruginosa oprF* interruption mutant produces twice as much biofilm as the parental strain (13). This result conflicts with a separate study in which an *oprF* mutant produced less biofilm when grown under anaerobic conditions (12). Furthermore, the OprF homolog OmpA, which is abundant in *Escherichia coli* biofilms (16), increases biofilm formation on hydrophobic surfaces (17). Mirroring this effect, in the pathogen *Acinetobacter baumannii, ompA* mutants are deficient in biofilm formation on abiotic surfaces and have decreased attachment to host cells (18). Together, these data suggest that OprF may play an important role in biofilm function.

Within the biofilm matrix, OprF is highly abundant in membrane vesicles, which are a major matrix component involved in biofilm structure and cell-to-cell signaling (5–19). Two membrane vesicle synthesis pathways have been established: the bilayer couple model, which produces outer membrane vesicles (OMVs), and the explosive cell lysis model, which results in membrane vesicles (20–21). Interestingly, OprF has been suggested to play a role in OMV production via the bilayer couple model. An OprF mutant overproduces OMVs relative to wild-type cells due to its overproduction of the quorum-sensing signal PQS (22). Since increased production of PQS and OMVs is correlated with biofilm dispersal (23), OprF may be important for this stage of the biofilm lifecycle. However, the role of vesicle-associated OprF in the biofilm is currently unknown (11).

Here we identified a nutrient-dependent biofilm defect in Δ*oprF* strains of *P. aeruginosa*. Upon dissection of the medium components, we found that Δ*oprF* biofilm formation was significantly reduced in the presence of glucose or low sodium chloride concentrations without affecting overall bacterial growth. The biofilm defect in the absence of OprF occurs during late-stage biofilm development and is not dependent on PQS production. Interestingly, we observed equivalent numbers of cells in mature wild-type biofilms and Δ*oprF* biofilms (that have reduced biofilm biomass). However, there was a significant reduction in eDNA in Δ*oprF* biofilms. Together, our data suggest that OprF is involved in the retention of eDNA during mature biofilm maintenance under certain growth conditions.

## RESULTS

### ΔoprF cells exhibit a nutrient-dependent biofilm defect

Since OprF is an abundant *P. aeruginosa* matrix protein (5–6), we tested the effect of deleting *oprF* on biofilm formation. We deleted *oprF* from *P. aeruginosa* PAO1 and confirmed via whole genome sequencing that our engineered deletion allele was the only difference between this strain and the parental strain. We also inserted an arabinose-inducible *oprF* at a neutral site in the chromosome in the Δ*oprF* background. This strain expressed OprF at levels similar to wild type upon addition of 0.5% arabinose, but not in the absence of inducer (Fig. S1). Using standard microtiter biofilm assays (24), we compared the Δ*oprF* biofilm formed in two common growth media: tryptic soy broth (TSB) and lysogeny broth (LB). While forming more biofilm than the exopolysaccharide-deficient Δ*pslD* negative-control strain in both media, Δ*oprF* formed 57.3 ± 3.8% S.D. (N=3; *p* < 0.01, ANOVA with post hoc Bonferroni) less biofilm than wild type in TSB, but an equivalent amount of biofilm to wild type in LB (Fig. 1A). This difference in biofilm formation was not due to growth rate differences in these media (see Fig. S2), and the biofilm defect was rescued in the inducible *oprF* strain when 0.5% arabinose was added. To determine if the Δ*oprF* biofilm defect in TSB exists in other *P. aeruginosa* strains, we constructed Δ*oprF* mutants in three other backgrounds: the tomato plant isolate E2, the water isolate MSH10, and the UTI isolate X24509 (25). Similar to PAO1, biofilm defects were observed in all three Δ*oprF* mutants when grown in TSB (Fig. S3A). Furthermore, the established *oprF* interruption mutant strain H636, which is made from a H103-based PAO1 background (26), exhibited a significant biofilm defect when grown in TSB (Fig. S3B). However, in agreement with a previously published study (13), the H636 strain produced approximately double the biofilm biomass as the H103 parental strain in LB (Fig. S3C). This H636 result conflicts with our Δ*oprF* strain biofilm phenotype in LB (Fig 1A), suggesting that the interruption mutation of *oprF* in H636 may be polar or that the H636 strain may have acquired secondary mutations. Nonetheless, since all Δ*oprF* strains that we tested had a biofilm defect in TSB, we continued our studies using our *P. aeruginosa* PAO1 Δ*oprF* strain.

**Figure 1.**
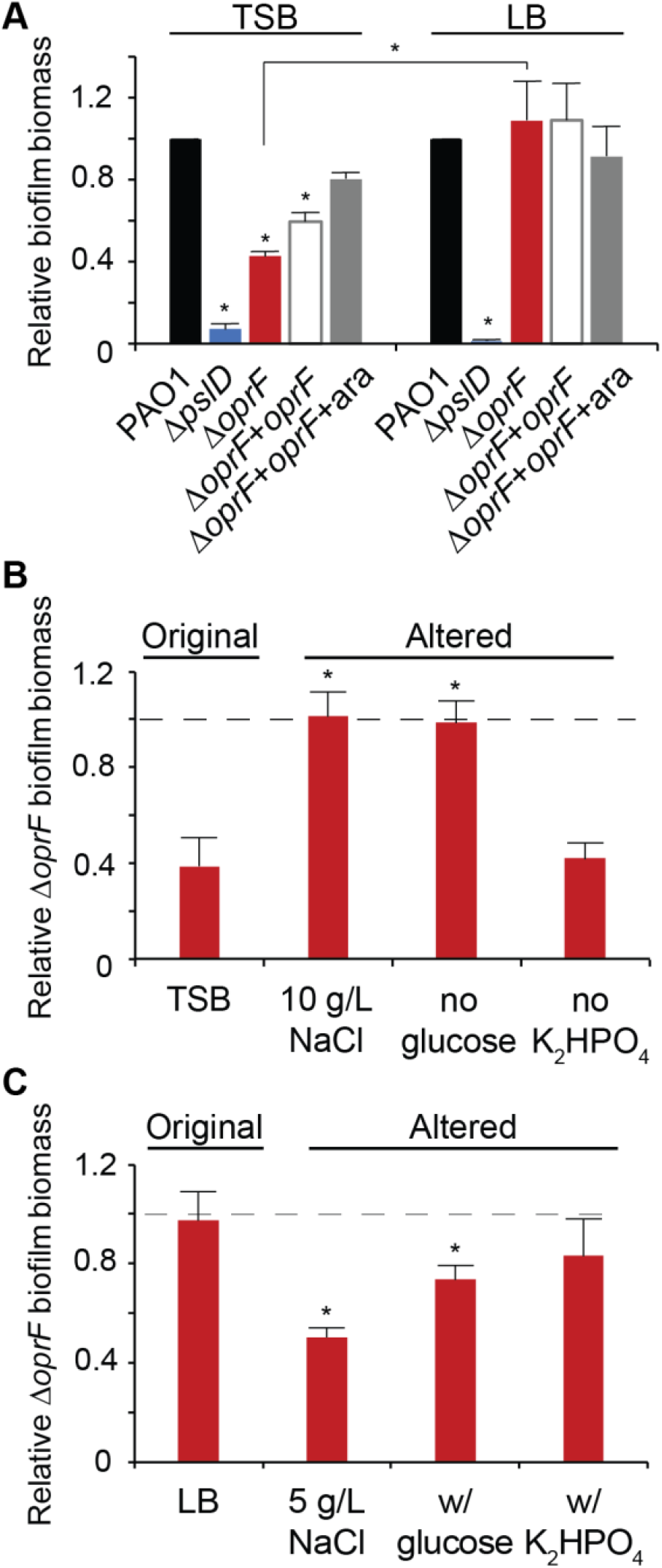
Δ*oprF* forms less biofilm in TSB than in LB, due to lower sodium chloride concentration and presence of glucose. (A) 24-hour static microtiter biofilm assays of *P. aeruginosa* PAO1 (WT, black), Δ*pslD* (blue), Δ*oprF* (red), and a Δ*oprF attTn7::P_BAD_-orpF* restoration strain (Δ*oprF* + *oprF*) without (white) and with (gray) 0.5% arabinose (ara) in the indicated media. Error bars, SEM (N = 3); asterisk over error bar, statistically different from WT in the same medium (p < 0.05; two-way ANOVA with post hoc Bonferroni). Statistical difference between Δ*oprF* strains in different media are indicated by a bar and asterisk. (B and C) Biofilm formation of Δ*oprF* strain in variations of TSB and LB: unaltered, altered NaCl concentrations, altered glucose concentrations, and altered K_2_HPO_4_ concentrations (left to right). Biofilm formation is normalized to WT in each respective medium. Dashed line, normalized amount of WT biofilm formation in each medium; error bars, SEM (N = 3); asterisk over error bar, statistically different from Δ*oprF* in the original medium (p < 0.05; two-way ANOVA with post hoc Bonferroni). See Figure S4 and Tables S2-S3 for full comparisons.

### Glucose and low sodium chloride reduce Δ*oprF* biofilm formation

While LB and TSB are both rich media with peptic digests as primary carbon sources, three notable ingredients differ between the two: sodium chloride (NaCl), glucose, and dipotassium phosphate (K_2_HPO_4_)(Table S1). To determine if these media components affect Δ*oprF* biofilm formation, we measured the static biofilm formed when strains were grown in media in which the concentrations of these components were individually altered to match that of the other medium. First, biofilms were grown in TSB or LB, each containing 5 or 10 g/L NaCl. Since reducing the NaCl concentration below 5 g/L decreases cell viability in *oprF* mutants (27), we did not test sodium chloride concentrations below this threshold. While Δ*oprF* formed less biofilm than wild type in TSB (with 5 g/L NaCl; original formula), Δ*oprF* formed biofilms similar to those of wild type when the NaCl concentration was increased to 10 g/L (with no other change in TSB) (Fig. 1B, S4A). The reciprocal effect was observed with LB, where Δ*oprF* formed biofilms similar to wild type in the original medium (with 10 g/L NaCl), but less biofilm than wild type when NaCl was reduced to 5 g/L (Fig. 1C, S4A). This reduced biofilm formation mirrors Δ*oprF* biofilms formed in TSB, which also contain 5 g/L NaCl. Changing the glucose concentration had a similar effect. Removing glucose from TSB resulted in Δ*oprF* biofilm biomass similar to that of wild type (Fig. 1B, S4B), mirroring the phenotype of Δ*oprF* biofilms formed in LB, which does not contain glucose (Fig. 1C, S4B). When glucose was added to LB, Δ*oprF* formed less biofilm than wild type, similar to biofilm formed by the mutant in TSB (which contains glucose). Changing the amount of K_2_HPO_4_ did not change the Δ*oprF* biofilm phenotype in either medium (Fig. 1B, 1C, S4C). These biofilm phenotypes were not the result of growth defects, as the planktonic growth rates of wild type and Δ*oprF* strains in these altered media were statistically equivalent (Fig. S2). These results indicate that Δ*oprF* biofilm formation is dependent on the NaCl and glucose concentrations.

### Δ*oprF* biofilm phenotype is not due to changes in osmolarity or metal concentrations

Since altering concentrations of major media solutes may impact medium osmolarity, we tested if the medium osmolarity is related to the Δ*oprF* biofilm defect by measuring the osmolarity of the various TSB and LB media with a vapor pressure osmometer and then correlating these measurements to the amounts of Δ*oprF* static biofilm biomass formed in the media. While there was a weak positive correlation between media osmolarity and Δ*oprF* biofilm formation, the relationship was not statistically significant (Fig. S5). We noted that the effect of osmolarity appeared to be driven by the changes in sodium chloride concentration within each medium (Fig. S5, squares). In comparison, glucose, which impacted Δ*oprF* biofilm formation, did not alter media osmolarity (Fig. S5, triangles). Assuming that the medium components impact biofilm formation through the same mechanism, we conclude that changes in osmolarity are not the major driving force behind the effect on Δ*oprF* biofilm formation.

Changes to media formulations can also affect the concentrations of biologically relevant metals. To determine the concentrations of iron, manganese, nickel, cobalt, copper, molybdenum, sodium, potassium, magnesium, calcium, and zinc, we performed inductively couple plasma mass spectrometry (ICP-MS) for each base medium and variant. While concentrations of sodium and potassium were altered when changes were made to sodium chloride or dipotassium phosphate levels, metal concentrations primarily tracked with TSB or LB base media (Fig. S6). Furthermore, there was no significant correlation between individual metal concentrations and Δ*oprF* biofilm formation (p > 0.05, Pearson’s). These results suggest that the nutrient-dependent effect of OprF in biofilm formation is not due to differential metal concentrations.

### OprF affects late-stage biofilm maturation in TSB

Biofilm formation occurs in distinct stages (28). The nutrient-dependent effects of OprF detailed above reflect mature static biofilm phenotypes and do not address when the Δ*oprF* biofilm defect in TSB begins. To pinpoint these potential time-dependent effects of OprF in biofilm formation, we performed static microtiter biofilm assays in TSB for 1,4, 8, 16, and 24 hours (29). There was no defect in the attached biomass of Δ*oprF* relative to that of wild type at any time point between 1-16 hours (Fig. 2). Unexpectedly, at 8 hours, Δ*oprF* formed more biofilm than wild type in TSB (Fig. 2C). However, by the 16-hour time point, Δ*oprF* biofilm levels once again matched those of wild type (Fig. 2D). These results suggest that the Δ*oprF* defect does not begin in the early stages of static biofilm formation. Instead, between the 16h and 24h time points, Δ*oprF* static biofilm biomass decreased by 36.8 ± 9.0% S.D. (N=3), while wild type increased 27.1 ± 16.1% S.D. (N=3). This suggests that without OprF, the static biofilm cannot maintain its biomass in TSB. Investigation of mature biofilm maintenance is ideally performed under continuous media flow, as it allows the biofilm to form for several days (30). However, the Δ*oprF* strain does not form biofilms on glass slides under media flow (unpublished data, E.K. Cassin and B.S. Tseng), limiting our methods to static assays. Combined with our earlier data on the nutrient-dependent effects of OprF, these data suggest that OprF is involved in the maintenance of mature *P. aeruginosa* biofilms in the presence of glucose or low sodium.

**Figure 2.**
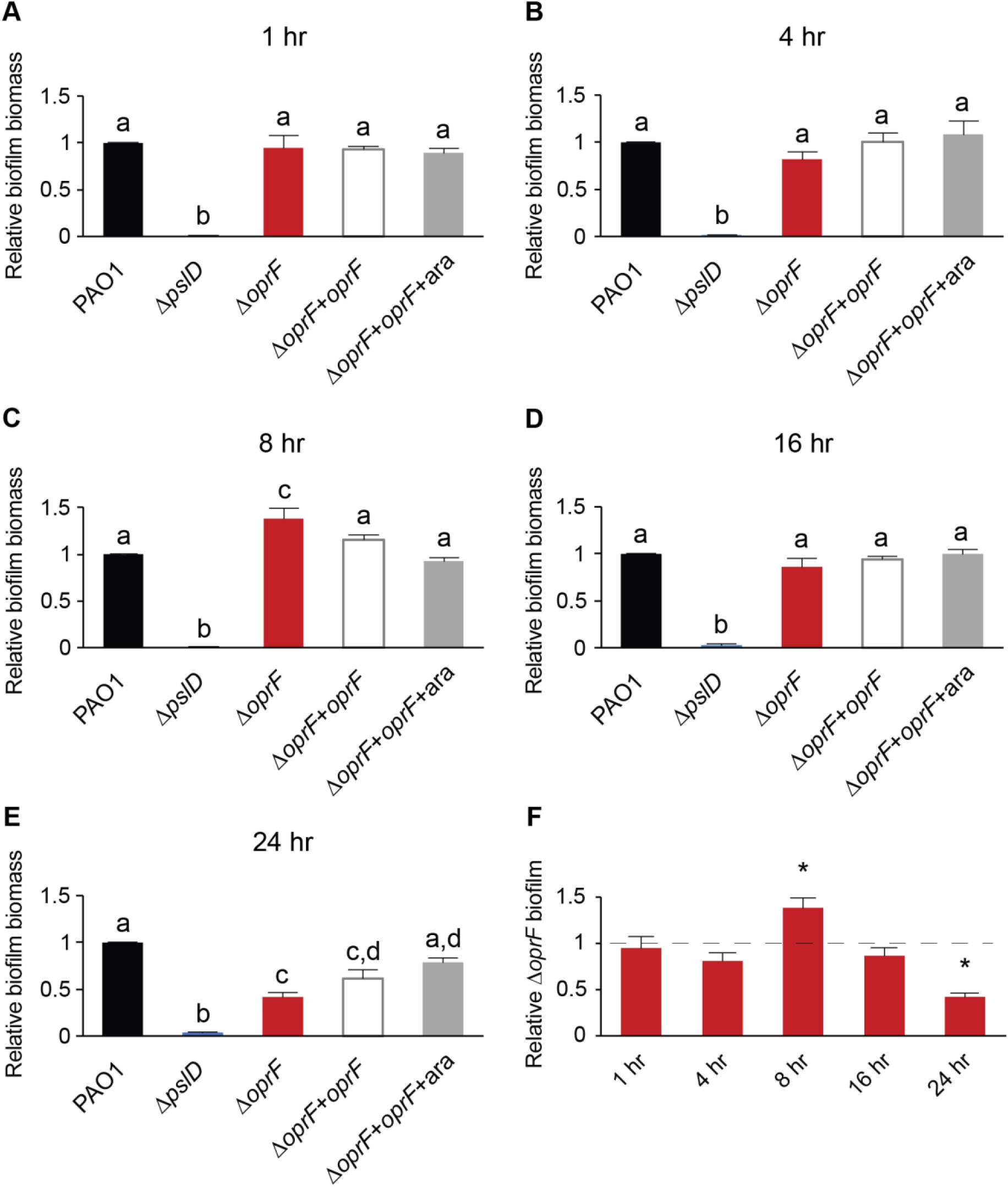
OprF affects late biofilm maturation in TSB. (A) 1-hour static microtiter biofilm assays were performed in TSB with PAO1 (WT, black), Δ*pslD* (blue), Δ*oprF* (red), and a Δ*oprF attTn7::P_BAD_-orpF* restoration strain (Δ*oprF* + *oprF*) with (white) and without (gray) 0.5% arabinose (ara). (B) 4-hour, (C) 8-hour, (D) 16-hour, and (E) 24-hour assays were performed with the same strains and media. (F) Static Δ*oprF* biofilm formation relative to WT (dashed line) at respective time points is represented. Biofilm formation is normalized to WT in each respective medium. Error bars, SEM (N = 3); letters, statistical groupings (p < 0.01; one-way ANOVA with post hoc Tukey HSD); asterisk, statistically different from WT at the same time point.

### The Δ*oprF* TSB biofilm defect is not dependent on PQS biosynthesis

The role of OprF in late-stage biofilm formation is interesting because planktonic *oprF* mutants make more OMVs than wild type cells and OMV production increases just before dispersal (22–23). We hypothesized that the Δ*oprF* biofilm defect in TSB may be due to an increased OMV production, resulting in early dispersion and less biofilm biomass relative to wild type. Since the increased OMV production of *oprF* mutants is due to PQS overproduction and deletion of PQS biosynthesis genes in an *oprF* mutant significantly decreases OMV production (22), we tested if deleting *pqsA* or *pqsH* in the Δ*oprF* strain would rescue the Δ*oprF* biofilm defect in TSB. Since PqsA is involved in the first steps of PQS biosynthesis and PqsH in the final step, a Δ*oprFΔpqsA* strain does not produce PQS or the PQS precursor HHQ, while a Δ*oprFΔpqsH* strain produces HHQ, but not PQS (31). While the Δ*pqsA* and Δ*pqsH* single deletion strains formed biofilms equal to wild type, both Δ*oprFΔpqsA* and Δ*oprFΔpqsH* formed biofilms equivalent to those of Δ*oprF* in TSB (Fig. 3), suggesting that increased PQS, and thereby OMV production, from the Δ*oprF* mutant strain is not responsible for the mature biofilm defect in TSB.

**Figure 3.**
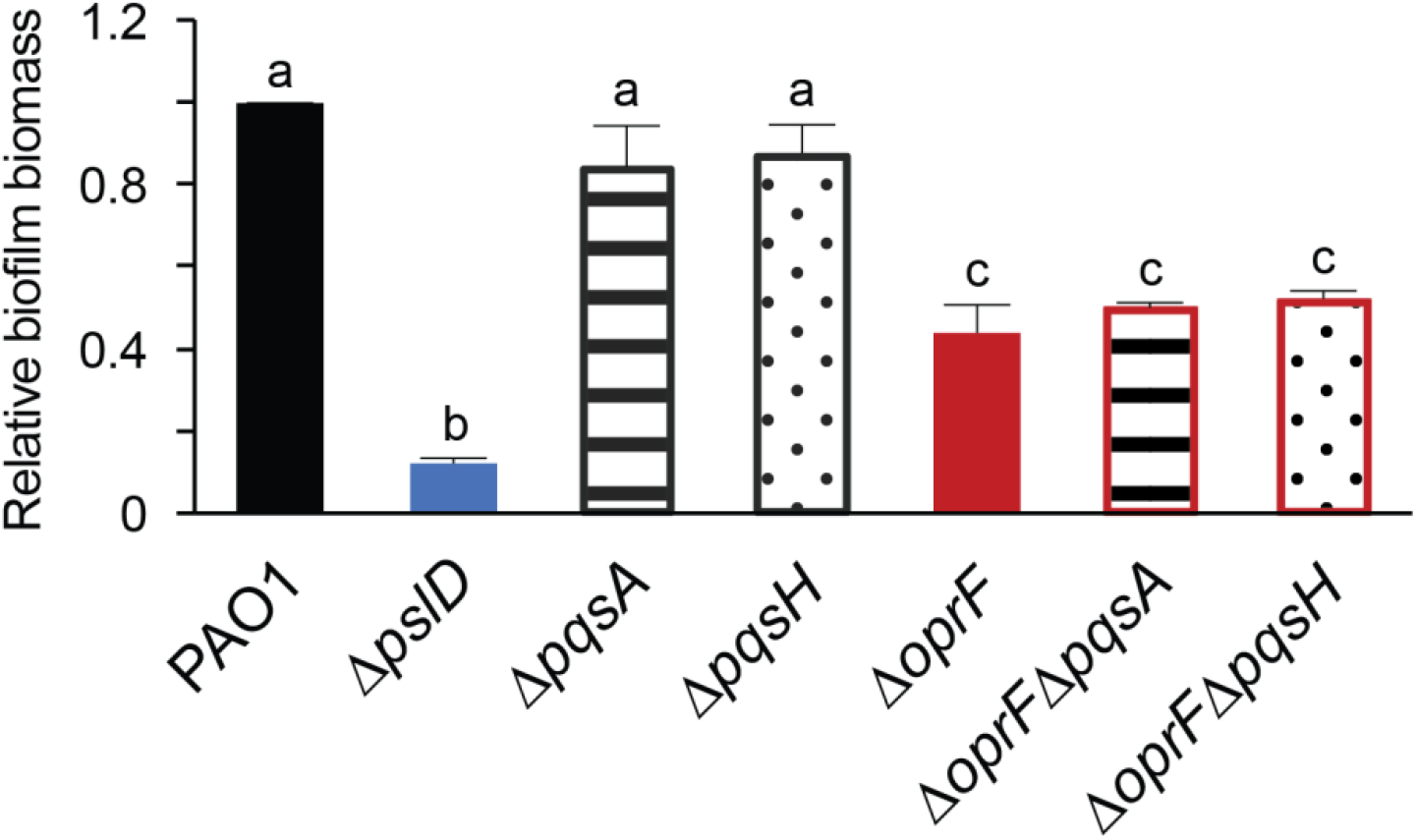
Δ*oprF* biofilm defect in TSB is independent of PQS. 24-hour static microtiter biofilm assays were performed in TSB with PAO1 (black), Δ*pslD* (blue), Δ*pqsA* (stripes), Δ*pqsH* (dots), Δ*oprF* (red), Δ*oprFΔpqsA* (stripes with red outline), and Δ*oprFΔpqsH* (dots with red outline). Biofilm formation is normalized to WT. Error bars, SEM (N = 3); letters, statistical groupings (p < 0.01; one-way ANOVA with post hoc Tukey HSD).

### Mature Δ*oprF* biofilms in TSB contain cell numbers equal to that of wild type

Static microtiter biofilm assays use crystal violet to stain surface-attached biomass as a proxy for total biofilm formation (29). Since crystal violet stains many biofilm components, including biofilm cells and the extracellular matrix, it is an indiscriminate indicator of surface-attached biomass. Therefore, we performed biofilm cell viability assays (29) in tandem with microtiter biofilm assays to tease apart which major components of the biofilm are affected by OprF. Surprisingly, despite the 60% decrease in total biofilm biomass in a side-by-side crystal violet staining (Fig. 4A), Δ*oprF* static microtiter biofilms in TSB contain approximately the same number of cells as that of wild type (Fig. 4B). Furthermore, to verify that the 60% decrease in Δ*oprF* static biofilms was not due to differential crystal violet staining between strains, we stained planktonic wild type and Δ*oprF* cells. These strains stain equivalently with crystal violet at the cell densities observed in the biofilm cell viability assays (Fig. S7). These results suggest that mature Δ*oprF* static biofilms in TSB contain less matrix, while biofilm cells remain attached to the surface and that OprF is involved in maintaining or retaining the mature biofilm matrix.

**Figure 4.**
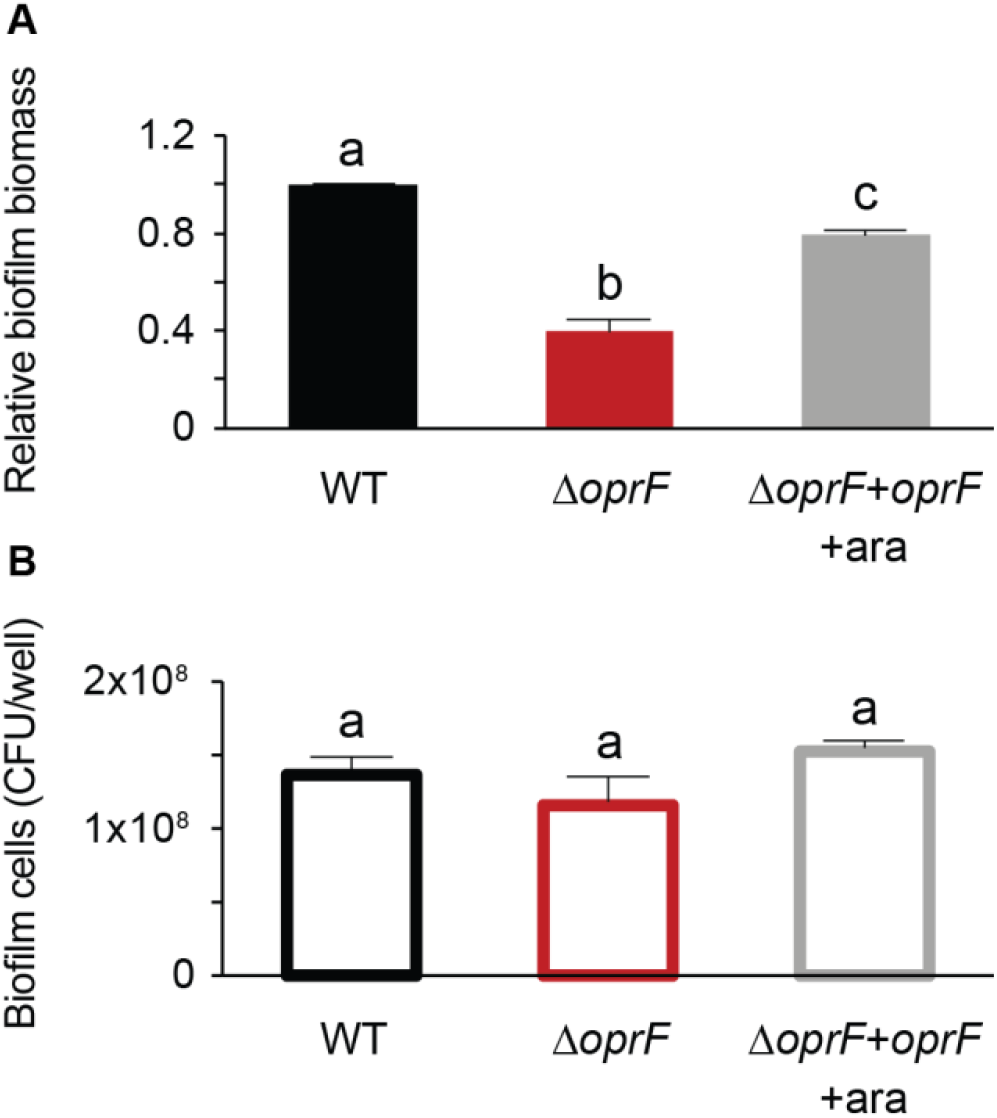
Δ*oprF* exhibits no biofilm cell viability defect in TSB. Side-by-side 24-hour (A) static microtiter biofilm and (B) biofilm cell viability assays were performed in TSB in the same 96-well plate with *P. aeruginosa* PAO1 (WT, black), Δ*oprF* (red), and the Δ*oprF att*Tn7::P_*BAD*_-*orpF* restoration strain with 0.5% arabinose (Δ*oprF* + *oprF* + ara; gray). Biofilm formation is normalized to WT. Error bars, SEM (N = 3); letters, statistical groupings (p < 0.05; one-way ANOVA with post hoc Tukey HSD).

### Δ*oprF* biofilms in TSB contain less eDNA than that of wild type

Since crystal violet stains negatively charged molecules, we reasoned that less eDNA in the biofilm could result in less biofilm biomass in the static biofilm assays. To quantify the eDNA in Δ*oprF* biofilms, we grew static Δ*oprF* or wild type biofilms in TSB and stained them with the eDNA-specific fluorophore DiTO-1. Static Δ*oprF* biofilms grown in TSB exhibit more eDNA-associated signal than the Δ*pslD* biofilm-negative control strain, but 58.6 ± 4.5% S.D. (N=3) less eDNA signal than wild-type biofilms (Fig. 5). This significant defect suggests that in the absence of OprF, eDNA is lost from the mature biofilm matrix. Furthermore, when combined with our earlier results, these results suggest that under certain conditions, OprF is involved in retaining eDNA in the mature *P. aeruginosa* biofilm matrix.

**Figure 5.**
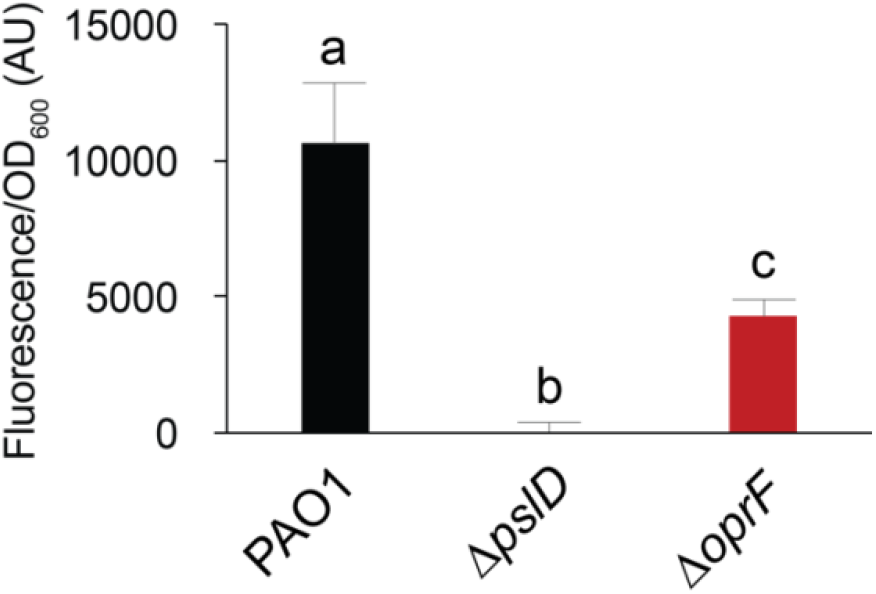
OprF affects biofilm eDNA levels in TSB. 24-hour static microtiter biofilms grown in TSB with PAO1 (black), Δ*pslD* (blue), and Δ*oprF* (red) were stained with the eDNA-specific dye DiTO-1. Fluorescence intensity from each strain was normalized to respective biofilm cell numbers (via absorbance at OD_600_). Error bars, SEM (N = 3); letters, statistical groupings (p < 0.05; one-way ANOVA with post hoc Tukey HSD).

## DISCUSSION

Our results highlight that growth conditions, specifically glucose and sodium concentrations, impact *P. aeruginosa oprF* mutant biofilm phenotypes. *P. aeruginosa ΔoprF* strains formed significantly less biofilm in TSB than LB. The decrease in Δ*oprF* biofilm in TSB occurred between 16-24 hours and did not result in fewer *P. aeruginosa* cells. Instead, Δ*oprF* biofilms in TSB contained significantly less eDNA than wild-type biofilms. The mechanisms underlying how glucose and low sodium led to decreased biofilms in cells lacking OprF is an exciting topic for future studies, as is determining how matrix-associated OprF affects eDNA levels.

Bouffartigues and colleagues previously found that an *oprF* interruption mutant forms approximately twice as much biofilm as the parental strain in LB, suggesting that a lack of OprF results in the overproduction of biofilm (13). Our results in LB using the same *oprF* interruption mutant strain agree with this conclusion. While these results follow the overall trend we saw in our Δ*oprF* strain in TSB and LB (Fig. 1), we did not observe hyperbiofilm formation in our Δ*oprF* strain in LB. Since both strains are of the PAO1 lineage and whole genome sequencing of our Δ*oprF* strain confirmed that no other differences exist between this strain and the parental, the difference in biofilm phenotypes suggests that there may be additional genetic factors at play. It is possible that the insertion in *oprF* in H636 affects biofilm formation or that the strain has accumulated secondary mutations within or outside the *oprF* interruption that affect biofilm formation in LB. These possibilities could be sorted out via future whole genome sequencing of H636 and comparing it to its parental strain.

Matrix-associated OprF, a membrane protein containing many hydrophobic residues, is abundant in biofilm membrane vesicles (4–5). OMV production in biofilms is dependent on PQS production (32), but in our experiments, abolishing PQS production did not impact the Δ*oprF* biofilm phenotype (Fig. 3). In a wild-type biofilm, cells produce OMVs via the bilayer couple model with PQS, and MVs via explosive cell lysis (32). In Δ*pqsA* biofilms, MVs are still produced (32), and we saw no defect in biofilm formation (Fig. 3). Similarly, MVs are likely still produced by cell lysis in the defective biofilms of both the Δ*oprF* and Δ*oprFΔpqs* strains. Notably, these mutant strains would produce vesicles with no OprF. Given that these strains exhibit 60% less biofilm than wild type, we conclude that this decline is due to the lack of OprF, independent of OMV production. Overall, the results of the current study indicate that in a Δ*oprF* background, PQS-mediated OMV synthesis is not related to the decrease in biofilm observed in TSB, which raises several questions outside the scope of this study: 1) do Δ*oprF* mutants in a biofilm produce more OMVs, as has been reported for planktonic *oprF* mutants (22)? 2) is matrix-associated OprF found only in vesicles? 3) how do glucose and low sodium affect the typical functions of OprF in biofilms? Further research probing these questions would expand our understanding of the roles of OprF and OmpA homologs in biofilm matrices.

OprF significantly affects the *P. aeruginosa* biofilm when grown under certain conditions. It is tempting to assume that the 60% decline in Δ*oprF* biofilms grown in TSB (Fig. 4A) is a proportional loss of all biofilm components. However, the static microtiter biofilm assay quantifies total biomass with crystal violet that stains the negatively charged components of the biofilm, namely cell surfaces, matrix membrane vesicles, and eDNA. Our biofilm cell viability assays demonstrate that Δ*oprF* biofilms do not lose 60% of their cells (Fig. 4B). Instead, the Δ*oprF* biofilms contain approximately 60% less eDNA than wild-type biofilms (Fig. 5). eDNA is an essential matrix component primarily produced by biofilm cell lysis (21–33). It has been proposed that membrane vesicles stabilize the matrix of wild-type biofilms through their interactions with eDNA (34). Therefore, OprF, which is abundant in membrane vesicles, may be involved – directly or indirectly – in these eDNA interactions and thereby in biofilm structural maintenance.

The maintenance of mature biofilms as an active, discrete stage in the biofilm lifecycle has been a recent topic of discussion (30). In this model, established biofilms respond to environmental changes to persist as a community. In a static microtiter biofilm, these changes include depletion of nutrients and waste accumulation over time. Our data indicate that OprF affects mature static biofilms in TSB, with the established Δ*oprF* biofilm decreasing between 16-24 hours of incubation. This phenotype suggests that in the absence of OprF, biofilm formation progresses to maturity and subsequently degrades. When combined with our biofilm cell viability results (Fig. 4), mature Δ*oprF* biofilm degradation does not appear to be due to dispersion since cell numbers are maintained. Therefore, we hypothesize that OprF may be involved in matrix retention in mature static biofilm maintenance via 1) matrix-bound OprF interactions with eDNA or 2) intracellular regulatory effects of deleting *oprF*. Future research into these lines of questioning is necessary and will contribute to an expanded understanding of the role of OprF in mature biofilm maintenance.

## METHODS AND MATERIALS

### Bacterial strains and growth conditions

Bacterial strains, oligonucleotides, and plasmids used in this study are in Tables S4-S6. Strains produced for this study were constructed using allelic exchange, as in (35), and described in Supplemental Methods. Liquid lysogeny broth (LB) and tryptic soy broth (TSB) were prepared according to the recipe in Table S1. The PAO1 Δ*oprF+oprF* strain containing *oprF* under an arabinose-inducible promoter was grown in media containing 0.5% L-arabinose (Sigma Aldrich). Unless otherwise noted, strains were grown at 37°C in specified media with 250 RPM shaking or on semi-solid LB containing 1.5% Bacto agar.

### Static microtiter biofilm assays

Static biofilms were grown as described in (24). Overnight cultures of bacteria grown in appropriate media were diluted 1:100, and 100 μL was seeded into sterile 96-well polystyrene plates (Greiner Bio-One, #650101). Plates were incubated at 37°C without shaking for 24 h unless otherwise noted. Planktonic cells were removed by triplicate washes in deionized water. Attached biofilm biomass was stained with 0.1% crystal violet for 15 min and washed as above. Stained biomass was solubilized using 30% acetic acid, transferred to a flat-bottom 96-well plate (Greiner Bio-One, #655090), and the absorbance at OD550 was read in a Synergy Hybrid HTX Microplate Reader (BioTek Instruments). Absorbance from blank media wells was subtracted from raw OD550 readings. Absorbance value of each strain was normalized to the average absorbance of the wild-type or parental strain. Four to ten technical replicates within each biological replicate were averaged, and the average measurement of three biological replicates were used to statistically compare biofilm formation by 1-way ANOVA with post hoc Tukey HSD for assays with 1 independent variable or 2-way ANOVA with post hoc Bonferroni for assays with 2 independent variables. All statistical analyses were performed in IBM SPSS.

### Biofilm cell viability assays

Biofilms were grown as above in static microtiter biofilm assays. Following 24-hr incubation, planktonic cells were removed by washing with sterile deionized water poured over plates three times. Half of the wells in each plate were scraped with sterile flat toothpicks in 125 μL sterile PBS to remove attached biofilm biomass. Solubilized biomass was serially diluted, spread on LB agar, and incubated at 37°C. CFU/well (100 μL/well) was enumerated after 24 h. The other half of the wells in each plate were stained with crystal violet, as detailed in static microtiter biofilm assays above. Four technical replicates within each biological replicate were averaged, and the average CFU/well of the three biological replicates was used to statistically compare cell counts by 1-way ANOVA with post hoc Tukey HSD.

### Biofilm eDNA fluorescence assays

Biofilms were grown as above in static microtiter biofilm assays. Following 24-hr incubation, planktonic cells were removed by washing with sterile deionized water poured over plates three times. Half of the wells in the plate were stained with eDNA-specific DiTO-1 (1 μM, AAT Bioquest, #17575) for 15 min. Stain was removed by pipetting and rinsed with 100 μL phosphate buffered saline in triplicate. Attached, stained biomass was removed by scraping with sterile toothpicks, as in biofilm cell viability assays above, in each well containing 125 μL sterile PBS. Scraped, stained biomass was transferred to a flat-bottomed, black-walled 96-well plate (Greiner Cellstar, #655090) and the fluorescence (Ex: 485/20, Em: 528/20) and absorbance (OD_600_) were measured in a Synergy Hybrid HTX Microplate Reader (BioTek Instruments). One quarter of the unstained wells were processed for cell viability and one quarter were processed for crystal violet staining to assess total biofilm formation, as above. The background fluorescent signal from wells incubated with media only was subtracted from total fluorescence, and the average total fluorescence from four technical replicates per biological replicate were averaged. The average fluorescence per biological replicate was normalized to the average OD_600_ value per strain. Average fluorescence/OD_600_ of the three biological replicates was used to statistically compare strain fluorescence by 1-way ANOVA with post hoc Tukey HSD.

## ACKNOWLEDGEMENTS

The authors would like to thank Kenesha Rae Broom, Mia G. Bruce, and Lindsey O’Neal for technical assistance, and Jeffrey W. Schertzer for critical discussions of this work. This project is funded by the NIH (K22 AI121097, P20 GM103440) and the Human Frontier Science Program (RGY00080/2021). In addition, EKC, DSB, and MCL were supported by NASA (80NSSC20M0043); SAA, DQR, and DSB, by NSF (#1301726, #1757316); and EKC and MCL, by University of Nevada Las Vegas Top Tier Doctoral Graduate Research Assistantships.

## AUTHOR CONTRIBUTIONS

Conceptualization: E.K. Cassin, B.S. Tseng; Formal analysis, investigation, methodology, or visualization of data: E.K. Cassin, S.A. Araujo-Hernandez, D.S. Baughn, M.C. Londono, D.Q. Rodriguez, B.S. Tseng; Writing – original draft: E.K. Cassin, B.S. Tseng; and Writing – review and editing: E.K. Cassin, M.C. Londono, B.S. Tseng.

